# Microtubule dynamics and the evolution of mitochondrial populations in fission yeast cells: A kinetic Monte Carlo study

**DOI:** 10.1101/2021.10.20.465075

**Authors:** Samlesh Choudhury, Vaishnavi Ananthanarayanan, K. Ganapathy Ayappa

**Affiliations:** Department of Chemical Engineering, Indian Institute of Science, Bangalore, Karnataka, India; EMBL Australia Node in Single Molecule Science, School of Medical Sciences, University of New South Wales, Australia

## Abstract

Mitochondrial populations in cells are maintained by cycles of fission and fusion events. Perturbation of this balance has been observed in several diseases such as cancer and neurodegeneration. In fission yeast cells, the association of mitochondria with microtubules inhibits mitochondrial fission, [1] illustrating the intricate coupling between mitochondria and the dynamic population of microtubules within the cell. In order to understand this coupling, we carried out kinetic Monte Carlo (KMC) simulations to predict the evolution of mitochondrial size distributions for different cases; wild-type cells, cells with short and long microtubules, and cells without microtubules. Comparison are made with mitochondrial distributions reported in experiments with fission yeast cells. Using experimentally determined mitochondrial fission and fusion frequencies, simulations implemented without the coupling of microtubule dynamics predicted an increase in the mean number of mitochondria, equilibrating within 50 s. The mitochondrial length distribution in these models also showed a higher occurrence of shorter mitochondria, implying a greater tendency for fission, similar to the scenario observed in the absence of microtubules and cells with short microtubules. Interestingly, this resulted in overestimating the mean number of mitochondria and underestimating mitochondrial lengths in cells with wild-type and long microtubules. However, coupling mitochondria’s fission and fusion events to the microtubule dynamics effectively captured the mitochondrial number and size distributions in wild-type and cells with long microtubules. Thus, the model provides greater physical insight into the temporal evolution of mitochondrial populations in different microtubule environments, allowing one to study both the short-time evolution as observed in the experiments (<5 minutes) as well as their transition towards a steady-state (>15 minutes). Our study illustrates the critical role of microtubules in mitochondrial dynamics and that coupling their growth and shrinkage dynamics is critical to predicting the evolution of mitochondrial populations within the cell.

**Author summary:** Mitochondria are semi-autonomous organelles that undergo fission and fusion to facilitate quality control and exchange of mitochondrial mass within the cell. Impaired mitochondrial fusion and fission dynamics are associated with disease states such as cancer and neurodegeneration. Recent experiments in fission yeast cells revealed a reduction in mitochondrial fission events when mitochondria were bound to the microtubules and longer microtubules shifted the mitochondrial population to longer lengths. In a distinct departure from earlier reports [2–16], we develop a generic framework to study the evolution of the mitochondrial population in fission yeast cells to predict the observed mitochondrial population by coupling the microtubule and mitochondrial dynamics. Using kinetic Monte Carlo (KMC) simulations we predict the temporal evolution of mitochondria in both the mutated and wild-type states of microtubules in fission yeast cells. The mitochondrial population evolves due to multiple fission and fusion reactions occurring between mitochondrial species of various lengths. Several models with varying complexity have been developed to study mitochondrial evolution, and predictions of the mitochondrial populations agree well with experimental data on fission yeast cells without microtubules and cells with short, wild-type and long microtubules. These set of microtubule states are consistent with not only the microtubule dynamics typically observed in cells under different physiological stimuli such as mitosis and disease states but also the stable microtubule states obtained through post-translational modification of *α* and *β* tubulin subunits of microtubules. Our study reveals that the temporal evolution of mitochondrial populations is an intrinsic function of the state of microtubules which modulates the fission and fusion frequencies to maintain mitochondrial homeostasis within cells.

## Introduction

Mitochondria, the primary sites of energy production in the cell are highly dynamic semiautonomous organelles [17–19]. Depending on the cell type and physiological stimulus, fission and fusion processes modulate both the mitochondrial numbers and morphologies. Mitochondrial fusion facilitates exchange of material (mitochondrial DNA, proteins, lipids and metabolites) between separate regions of the mitochondrial network, thus allowing the repair of defective mitochondria and efficient utilisation of available substrates [20]. Fission on the other hand plays an important role in quality control of mitochondria by mitophagy [21]. Thus the interplay of fission and fusion processes is responsible for maintaining a functional mitochondrial population [22, 23]. Perturbations in the fission-fusion dynamics of mitochondria is closely related to neurodegenerative diseases such as Alzheimer’s, Huntington’s, or Parkinson’s [24–29]. A variety of mathematical models have been used to study mitochondrial dynamics [2–7] to establish the role of fission and fusion processes in regulating mitophaghy [30] pathways, ageing related accumulation of mitochondrial DNA mutations [31, 32] and maintaining the necessary bioenergetic supply and demand cycle of the cell [3]. These studies have emphasized the impact of mitochondrial locomotion along microtubule tracks on the fission and fusion phenomenon, however the inherent dynamic nature of microtubules and their coupling with the mitochondrial population redistribution is incompletely understood. In contrast, the dynamics of microtubules in the absence of mitochondria has been extensively investigated in the literature.

Microtubules are hollow cylindrical structures consisting of 13 protofilaments made of *α* and *β* tubulin heterodimers. Each *α* and *β* tubulin heterodimer is 8nm long, and the overall diameter of the microtubule is approximately 25 *µ*m [33]. The polarity of microtubules arises from the head-to-tail arrangement of *α* and *β* tubulin dimers in each of the protofilaments. Such polarity in microtubule structures is central to the inherent polymerization and depolymerization dynamics of microtubules. During polymerization, the *β* tubulin undergoes GTP hydrolysis, and the resulting GDP-bound structures undergo catastrophe events when the rate of hydrolysis exceeds the rate of polymerization of the microtubule. During depolymerization, the addition of GTP-bound tubulin subunits to the plus end of the protofilaments leads to a rescue event. The stochastic switching between polymerization and depolymerization is known as the dynamic instability of microtubules. Microtubule dynamics is essential for exploration of cellular space, trafficking of poorly diffusive cargo in the cell and also self organising properties of the mitotic spindle [34–39]. Several analytical models have been developed to understand microtubule behaviour through both deterministic and stochastic methods such as Monte Carlo approaches, providing insight into the inherent instabilities associated with microtubule dynamics [8–12, 14, 16]. The dynamic coupling of mitochondrial fission and fusion events with the inherent stochastic and evolving population of microtubules within the cell has yet to be investigated. This intricate coupling has been shown to play an important role in maintaining mitochondrial homeostasis in fission yeast cells [1].

In these experiments by Mehta et al., [1, 40], a reduction in fission events was observed when mitochondria were bound to microtubules. Fission yeast mitochondria bind to microtubules through the Mmb1p protein [41] and do not undergo motor driven movement along the microtubules [42, 43]. Cells with long microtubules had longer, but fewer mitochondria, whereas cells with short microtubules had shorter, but several more mitochondria than wild-type (WT) cells. These experiments reveal the dynamic coupling between mitochondria and microtubules in determining both the evolution of the mitochondrial numbers and size distributions within the cell. With this perspective, the primary focus of this manuscript is to develop a suitable model to study the evolution of the mitochondrial population by including the coupling with the microtubule dynamics within the cell.

We employed kinetic Monte Carlo (KMC) [44–46] simulations to model the evolution of mitochondrial population in cells in the absence of microtubules (MT_absent_) as well as in the presence of short (MT_short_), long (MT_long_) and WT microtubules(MT_wt_) and use experimental data from the study of Mehta et al., [1], to compare the predicted mitochondrial numbers and sizes. The mitochondrial population was assumed to be an outcome of multiple fission and fusion events occurring between mitochondrial species of various sizes while conserving the overall mass of the mitochondria for the time scales under consideration. In this modeling effort, the major challenges faced in quantifying the dynamics were posed by the low mitochondrial population number and the high degree of stochasticity associated with their evolution. In the first approach (model M1), the fission yeast cell was assumed to be a well mixed system and the mitochondria population evolves using a Gillespie algorithm with experimentally defined fission and fusion rate constants. In the second approach, the mitochondrial population was evolved in a biologically more relevant environment similar to the experiments in fission yeast cells [1]. In experiments, mitochondria were observed to be largely static lacking the freedom to diffuse across the entire cellular volume and fission events were suppressed when mitochondria were associated with microtubules. To emulate these conditions, we carried out simulations where the cellular volume was halved along the width of the cell and mitochondria populations were evolved in these newly remodelled cellular volumes called sectors. In this model, simulations were carried out both in the absence (model M2) and presence (model M2MT) of microtubules . In the M2MT model, the dynamics of different classes of microtubules was obtained independently and subsequently coupled to the mitochondrial population dynamics. Although, models implemented without coupling to the microtubule dynamics (M1, M2) captured the mitochondrial evolution in MT_absent_ and MT_short_ cells, incorporating the coupling with the microtubule dynamics enabled us to capture the mitochondrial dynamics similar to those observed in the experiments by Mehta et al., [1] under different conditions of microtubule growth.

## Kinetic Monte Carlo Simulations

### Distribution of Mitochondrial Lengths

For a given state of the system, mitochondria were treated as linear fragments undergoing fission and fusion events. From experimental data on fission yeast cells [1] the shortest and longest lengths observed across the different mitochondrial populations were *L*_*min*_ = 0.6 *µ*m and *L*_*max*_ = 12.6 *µ*m respectively. Based on *L*_*min*_ and the ratio of *L*_*max*_ to *L*_*min*_, 20 different length fragments were constructed and the experimentally observed mitochondrial lengths were binned to create a histogram yielding lengths between *L*_*i*_ and *L*_*i*+1_, denoted by *N*_*i*_(*t*) to represent the number of mitochondria present in fragment *i* at time *t*. Further, *N* (*t*) ≡ [*N*_1_(*t*), *N*_2_(*t*), …, *N*_20_(*t*)] defines the overall state of the system at a particular time.

### Reaction Schemes

Mitochondrial species evolve through a network of fission and fusion reactions. Popular mass action law based models such as the Smoluchowski coagulation-fragmentation kinetics [47–50] were used to represent fission and fusion reaction rates. Similar to a general binary Smoluchowski coagulation-fragmentation process, fission processes were assumed to follow first order reaction kinetics whereas fusion reactions were considered to be second order. The rate constants *K*_*fission*_ and *K*_*fusion*_ used in these reactions were derived from the experiments on fission yeast cells [1]. Hence, the rates for a fission reaction (*r*_*fission*_) of species *i* and a fusion reaction(*r*_*fusion*_) between species *i* and *j* were defined as follows:

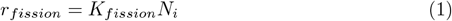

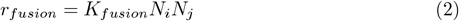

We further assume there is neither mitochondrial biogenesis nor mitophagy for the time scales under consideration.

### Gillespie Algorithm

The direct Gillespie algorithm [51, 52] was chosen as the kinetic Monte Carlo (KMC) scheme to capture the dynamics of the mitochondrial system. In the simulations, based on the binning outcome of the 20 different length fragments, a fission-fusion reaction network consisting of 201 reactions was initiated. Evolution in the number of mitochondria present in each fragment can occur in two ways: i) fusion reactions with mitochondria from neighboring fragments ii) fission reactions of the mitochondria present in the current fragment. Mitochondrial lengths were obtained from experiments by Mehta et al., [1] carried out in 14 MT_absent_ cells, 8 MT_short_ cells, 15 MT_wt_ cells and 21 MT_long_ cells. All experiments correspond to a time period of 228 seconds. The lengths were then partitioned into predefined mitochondria fragments as per the above binning procedure and number of mitochondria present in individual fragments were obtained. Thus for each cell the number of mitochondria present in each fragment at time *t* = 0 defines the initial state *N* (*t*_0_) of the cell. The Gillespie algorithm was then carried out using *N* (*t*_0_) as the initial condition and the time evolution of the state vector of respective cells was obtained from the model.

## Modeling Approaches

We outline various modelling strategies that were used to simulate the temporal evolution of mitochondria with the KMC framework. In the first approach (model M1), the entire fission yeast cell was assumed to be a well mixed system (Fig 1A) and the mitochondria population was evolved using the Gillespie algorithm with the experimentally defined fission-fusion rate constants. In the second approach, which we refer to as the M2 models, the cellular volume was halved along the width of the cell (Fig 1B and Fig 1D) to create sectors and the mitochondria populations were independently evolved in each sector. We refer to simulations carried out in the absence of microtubules as M2 model (Fig 1B) and simulations in the presence of microtubules as the M2MT model (Fig 1D). The microtubule dynamics incorporated into the M2MT model was obtained independently using a four sector partitioning of the cell (Fig 1C). Flowcharts for the M1, M2 and M2MT models are given in S1 Fig., S2 Fig. and S4 Fig. respectively.

**Fig 1.**
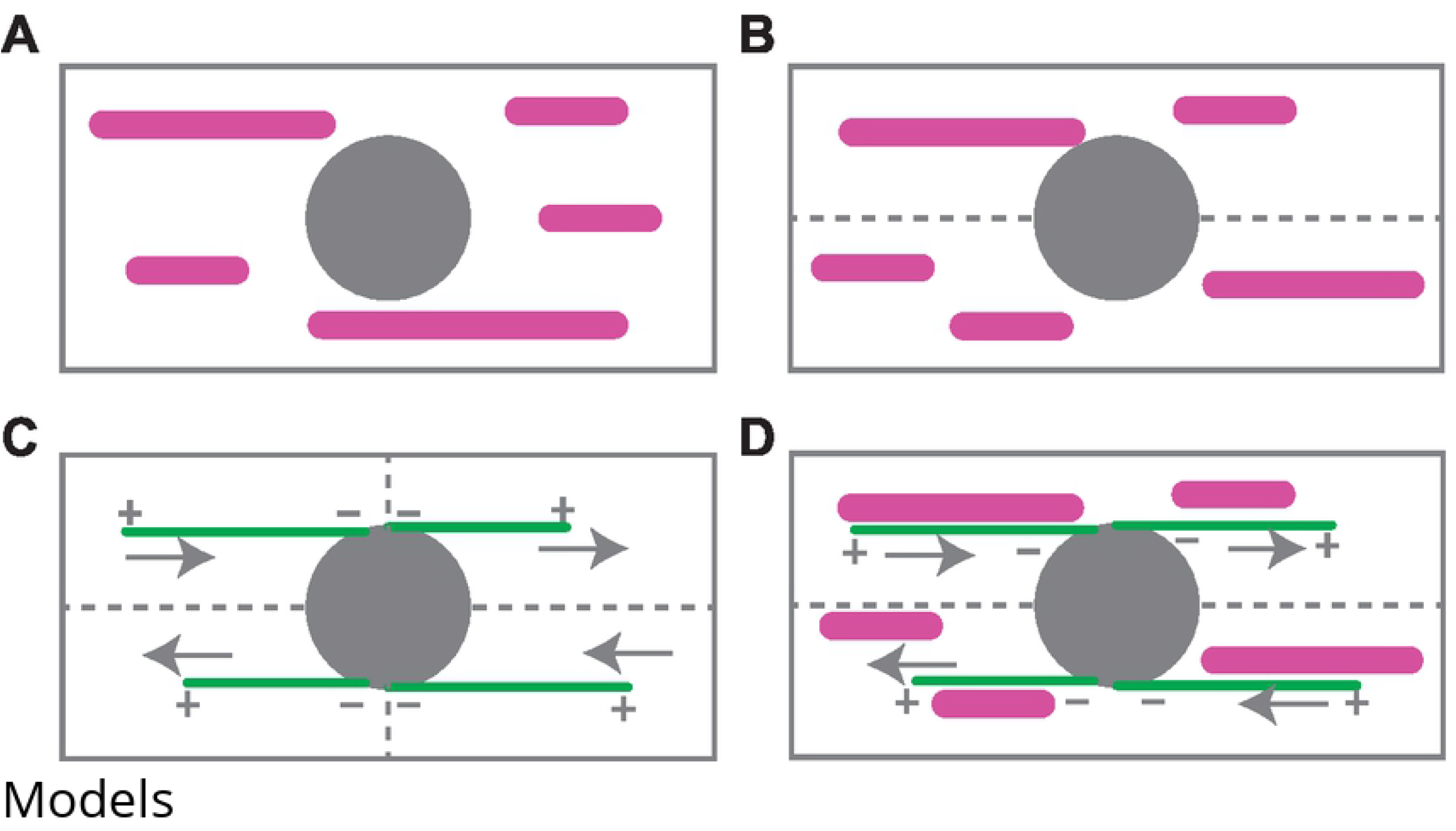
Mitochondria-Microtubule models. (A) M1 model for the yeast cell with the nucleus (grey) centrally located and mitochondria (pink) distributed throughout the cell.(B) M2 model wherein the cell is partitioned (grey dotted lines) into sectors and mitochondria are distributed randomly in the two sectors (C) Yeast cell is divided in four sectors and in each sector a microtubule bundle (green solid lines) is evolved to obtain microtubule dynamics as represented by grey arrows. (D) M2MT model in which two sectors are present in order to simulate the dynamics of mitochondria in the presence of microtubules

### Microtubule Dynamics

Microtubules were modeled as linear polymers comprising of heterodimeric subunits of *α* and *β* tubulin. The *β* tubulin also possesses an exchangeable GTP molecule. As a result, in a hydrolysis reaction where GTP converts to GDP, the subunits undergo a steric change. Hydrolysis and polymerization processes were independent of each other. During polymerization, the microtubules grow outward from the microtubule-organising centre with GTP-bound [53] subunits and progress towards the cell wall along the long axis of the cell. Hydrolysis of other tubulin subunits in the polymer also occur simultaneously [8]. If the rate of hydrolysis of subunits exceeds the rate of polymerization then microtubules switch to a state of depolymerization, in an event referred to as catastrophe. However, if the microtubules grow to the cell wall, a pause event occurs and hydrolysis precedes complete depolymerization [54].

Polymerization reaction propensities (*P*_*poly*_) follow first order reaction kinetics and is dependent on soluble tubulin concentration (*C*_*T*_) present in the cytoplasm [8]. The propensities of hydrolysis (*P*_*hy*_) and depolymerization (*P*_*depoly*_) follow zero order kinetics. The initial concentration of tubulin was assumed to be 3 *µ*M [55]. Thus, the overall dynamics of microtubules follow the associated propensities, *P* as follows:

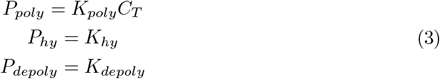

Yeast cells are typically 8 − 13 *µ*m long [56] in the interphase stage and we assumed a cell length of 11.6 *µ*m in all the simulation models. To simulate microtubule dynamics, the yeast cell was equally divided into four virtual sectors as shown in Fig 1C such that the length of each sector was 5.8 *µ*m. In each sector, only one microtubule bundle was simulated similar to the arrangement of microtubules observed in experiments [55]. Every microtubule bundle consisted of three microtubules, as noted in experiments [55] and the microtubule plus ends were towards the periphery as shown in Fig 1C and Fig 1D. In a polymerization event, the microtubule bundle length increased such that the centre of mass of the bundle was displaced linearly by 0.1 *µ*m as shown in Fig 1C. However, when a hydrolysis event was triggered, all the subunits in the particular cross-section of 0.1 *µ*m width underwent hydrolysis. Thus, in a sector, when all the subunits were hydrolysed, a catastrophe event occurrs and the length reduces to zero. The Gillespie algorithm was implemented to simulate different microtubule environments based on the microtubule kinetics given in Eq 3. Using the simulated lengths of microtubules, various microtubule parameters such as elongation velocity 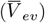, shrinkage velocity 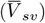 and catastrophe frequency 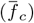 [8] were computed and compared with experiments. The details of these computations are in S5 Text. The set of rate constants [*K*_*poly*_, *K*_*hy*_, *K*_*depoly*_] for polymerization, hydrolysis and depolymerization respectively, used in Eq 3 were obtained by applying a constrained optimisation genetic algorithm [57], using the routine ‘ga’ in Matlab2019b to minimise the objective function defined as the difference between the simulation and experimental values reported for the microtubule parameters 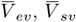 and 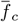.

To evolve the mitochondrial populations in model M2MT, the microtubule length data obtained from the pair of horizontally adjacent sectors as present in Fig 1C were summed up at common time interval to obtain the temporal microtubule length distribution for the newly formed pair of vertically adjacent sectors to be used in model M2MT (Fig 1D). Initially the mitochondria present in the state vector *N* (*t*_0_) were randomly distributed among the two virtual sectors along the width of the cell (Fig 1D). At a given instant in time, from the state vector *N* (*t*), the lengths of mitochondria present in a particular sector were identified and compared with the lengths of the microtubule present in that particular sector. Mitochondria with lengths less than the microtubule were allowed to undergo only fusion reactions. However, for mitochondria having lengths greater than that of the microtubule, the fraction of the length of the mitochondria greater than that of the microtubule could undergo both fission and fusion reactions. The rest of the mitochondria could undergo only fusion reactions.

## Results and Discussions

To interface with the available experimental data [1], KMC simulations were carried out in 14 MT_absent_ cells, 8 MT_short_ cells, 15 MT_wt_ cells and 21 MT_long_ cells. Each stochastic simulation was evolved for at least 500 s and 1000 replicates of a particular simulation were implemented for every cell across the different microtubule environments. In every cell with different microtubule environments, the experimental length data were binned and the number of mitochondria corresponding to the initial state observed in the experiment was used as the initial condition for the simulations. The rate constants of fission and fusion used in each of these stochastic simulations are tabulated in Table 1. Hence at a particular time step, the overall mean number of mitochondria, 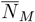 for a specific microtubule environment was obtained by averaging the number of mitochondria across the 1000 replicates and all the cells for a particular microtubule environment.

**Table 1.**
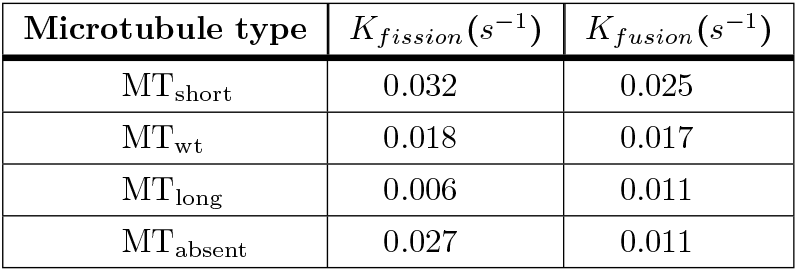
Mitochondria fission and fusion rate constants in fission yeast cells with different microtubule environments [1].

### Constraint free mitochondrial evolution and effect of sector constraints on their evolution

We first studied the temporal evolution of mitochondria species for the models M1 and M2. In both these models, an increasing trend in the mean number of mitochondria, 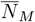 was observed at the initial time for MT_absent_ cells (Fig 2A) and MT_short_ cells (Fig 3A). The time taken to reach an equilibrium value was about 40 s in both the models M1 and M2, however a more gradual evolution of the mitochondria population was observed in the experiments, appearing to saturate above 200 s. For Mt_short_ cells (Fig 3A), the dynamics was faster in the experiments with a rapid saturation in the mean mitochondria number observed between 40-50 s. Interestingly model M2 was able to capture the equilibrium population of mean mitochondrial numbers more accurately when compared with M1, at longer times (Fig 3A). The variation in the mitochondrial length distributions, *L*_*M*_ illustrated in Fig 2B and Fig 3B yield greater insight into the evolution of the mitochondrial sizes for the MT_absent_ and MT_short_ cells respectively. For both cases we observed that the mean and the spread of mitochondrial lengths for model M2 agreed well with the experiments, however for model M1 the mean was shifted to smaller mitochondrial lengths with reduced spread as observed in Fig 2B and Fig 3B. In the case of model M1, the lower mean length was consistent with the larger *N*_*M*_ observed in the time evolution data (Fig 2A and Fig 3A).

**Fig 2.**
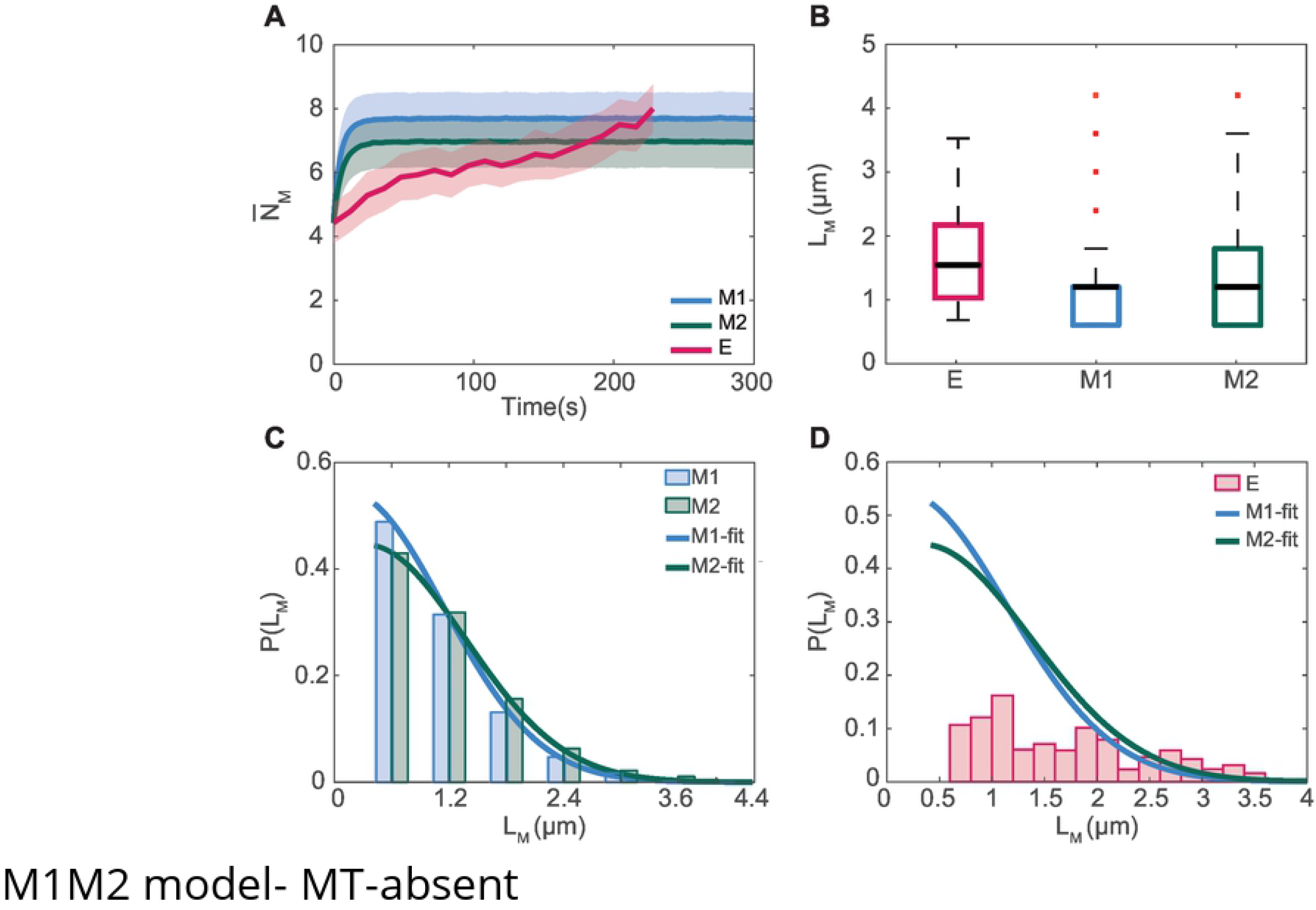
Comparison of the number of mitochondria, 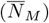 and distributions of mitochondria lengths, *L*_*M*_ obtained from the models M1 and M2 with experimental data (E) for the MT_absent_ cells. (A) Temporal evolution of 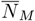 . (B) Comparison of *L*_*M*_ .(C) Probability distribution, P(*L*_*M*_) of mitochondrial lengths in simulations with the Gaussian fits and comparison with experiments (D)

**Fig 3.**
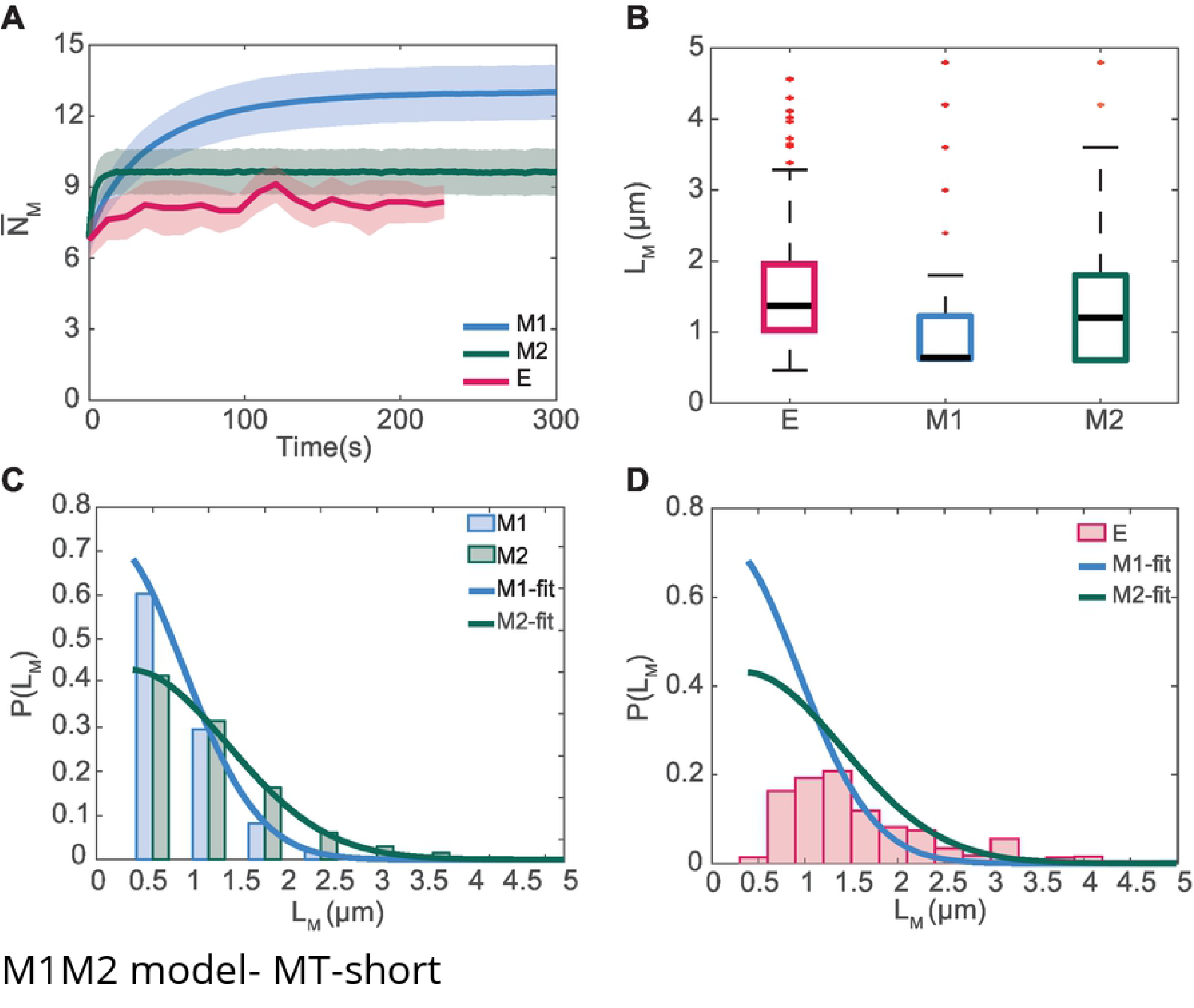
Comparison of the number of mitochondria, 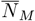 and distributions of mitochondria lengths, *L*_*M*_ obtained from the models M1 and M2 with experimental data (E) for the MT_short_ cells. (A) Temporal evolution of 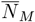 . (B) Comparison of *L*_*M*_ . (C) Probability distribution, P(*L*_*M*_) of mitochondrial lengths in simulations with the Gaussian fits and comparison with experiments (D)

The probability distribution of mitochondria lengths (P(*L*_*M*_)) predicted from both the M1 and M2 models peaked around the shortest mitochondria lengths of 0.6 *µ*m as illustrated in (Fig 2C and Fig 3C). A Gaussian distribution was seen to accurately capture the length variation obtained from both the M1 and M2 modes. We superimposed these fits on the experimentally observed length distributions (Fig 2D and Fig 3D) which revealed that both models overestimated the population at smaller lengths. The mitochondrial length distributions obtained in experiments were generally broader with a greater population of longer mitochondria (Fig 2D and Fig 3D). This trend was qualitatively captured to a better extent with the model M2 for MT_short_ as illustrated in Fig 3D. Hence, although both models M1 and M2 qualitatively captured the observed trends in mean numbers of the mitochondrial population, model M2 provides improved predictions from both the mean mitochondrial numbers as well as the lengths when compared with the experiments.

We next discuss results for the Mt_wt_ and MT_long_ environments. We point out that in the experiments, the mean number in the mitochondrial population 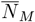 remain largely constant for the MT_wt_ (Fig 4A) and higher than that of MT_long_ (Fig 5A) since the mitochondrial fission and fusion frequencies were comparatively higher in MT_wt_ cells [1]. The predictions of the mean number of mitochondria from both model M1 and M2 models were considerably higher, overestimating the numbers by about a factor 2. Interestingly we observed the numbers at equilibrium to be larger for M2 when compared with M1. This was in contrast to the earlier cases in the MT_absent_ and MT_short_ cells where opposite trends were observed (Fig 2A and Fig 3A). The mitochondrial length distributions predicted by both models M1 and M2 models were shifted toward shorter lengths when compared with the observations in experiments (Fig 4B and Fig 5B). The mitochondrial lengths for MT_wt_ and MT_long_ cells (Fig 4B and Fig 5B) were also higher when compared with those in MT_absent_ cells and MT_short_ environments (Fig 3B and Fig 2B). The increased propensity toward shorter mitochondria in the mitochondrial population predicted from the KMC simulations leads to a greater difference from the mitochondrial populations observed in experiments for both MT_wt_ and MT_long_ cells (Fig 4B and Fig 5B). Similar to the trends observed for MT_absent_ cells and MT_short_ cells, the mitochondria length distributions (P(*L*_*M*_)) predicted from both the M1 and M2 models were peaked around the shortest mitochondria lengths as illustrated in (Fig 4C and Fig 5C). The deviation from experimental data was specifically stark for these situations with the greatest differences observed for the MT_long_ where mitochondrial lengths are severely underestimated (Fig 5D).

**Fig 4.**
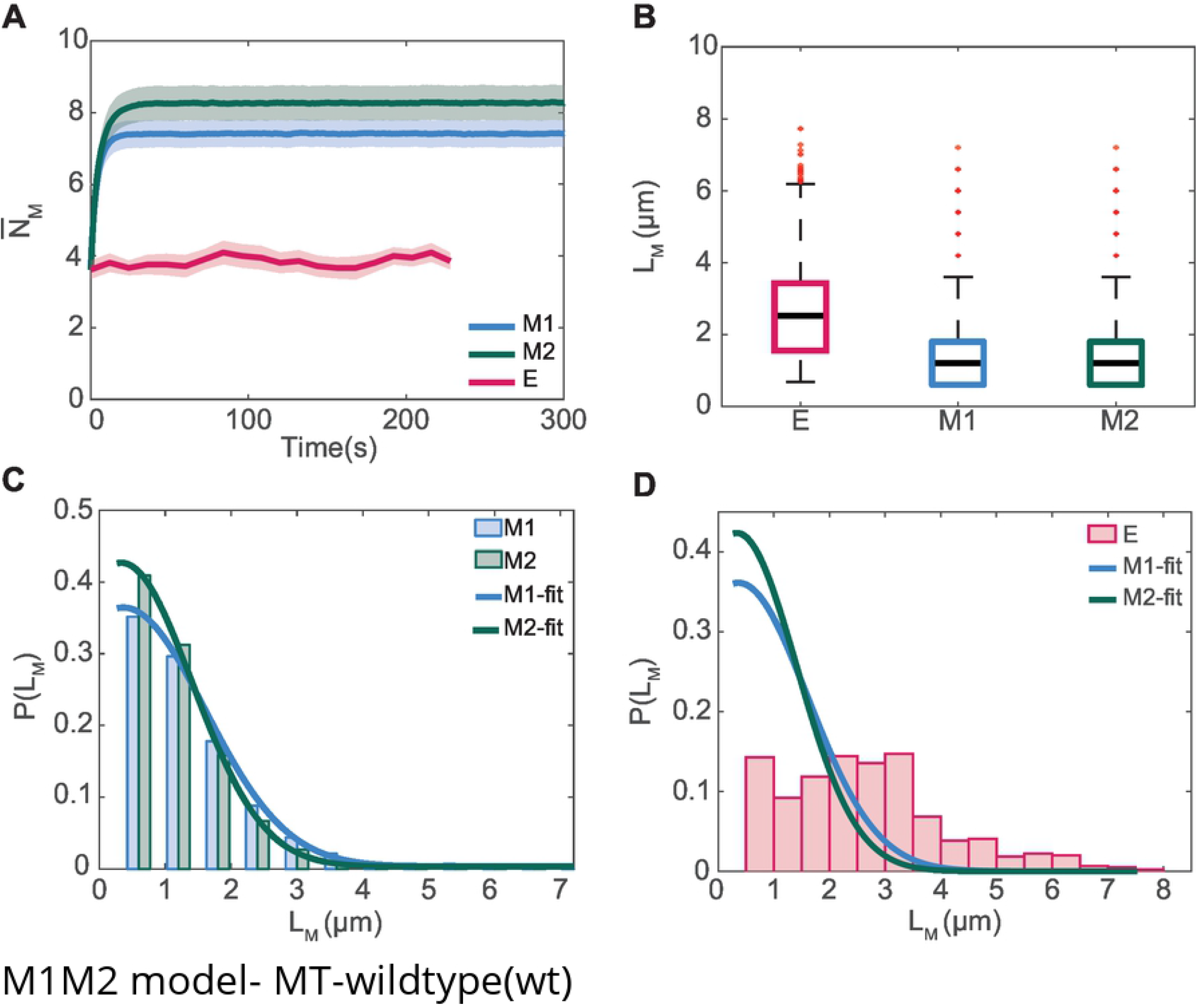
Comparison of the number of mitochondria, 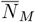 sand distributions of mitochondria lengths, *L*_*M*_ obtained from the models M1 and M2 with experimental data (E) for MT_wt_ cells. (A) Temporal evolution of 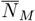. (B) Comparison of *L*_*M*_ . (C) Probability distribution, P(*L*_*M*_) of mitochondrial lengths in simulations with the Gaussian fits and comparison with experiments (D)

**Fig 5.**
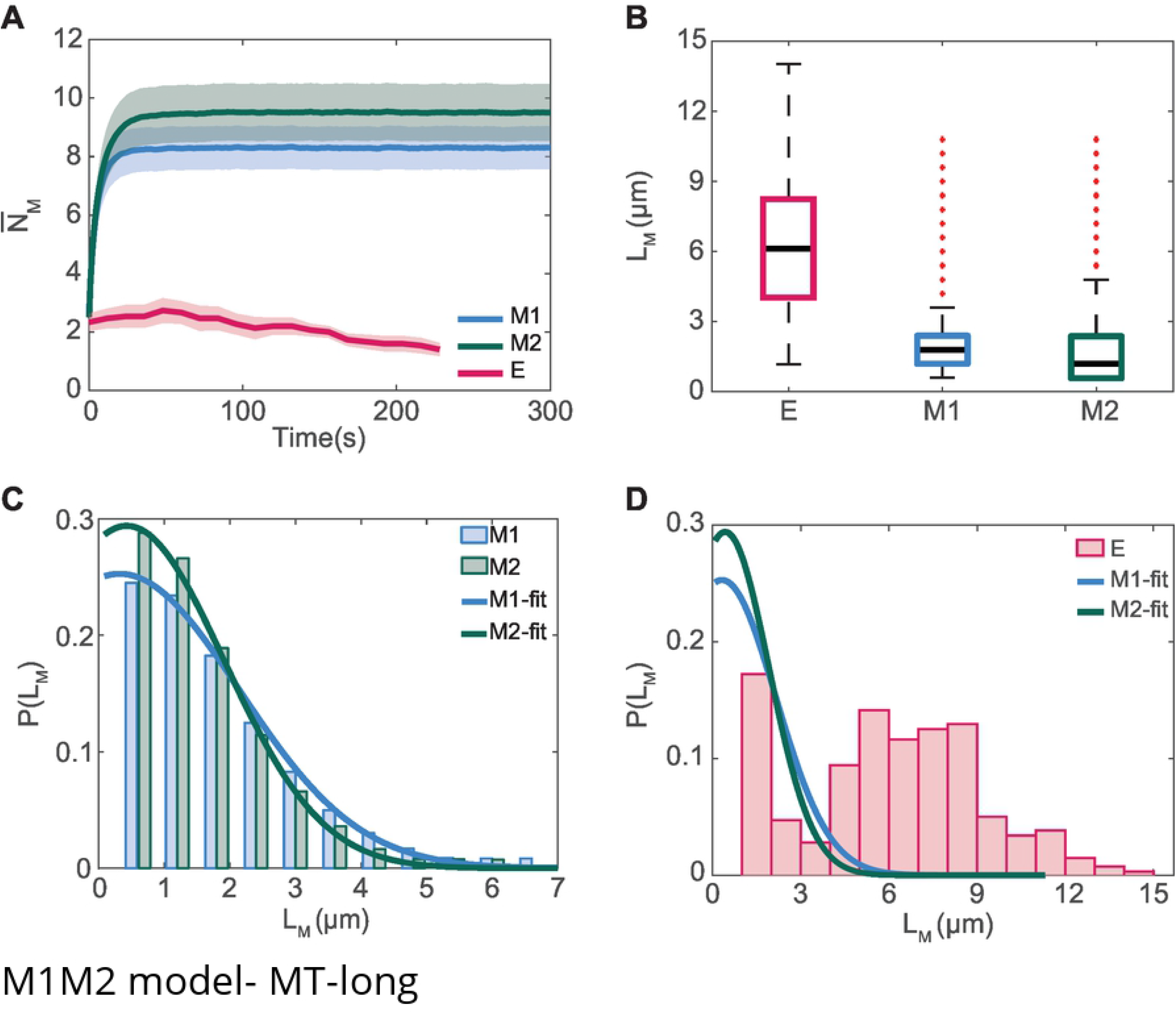
Comparison of the number of mitochondria, 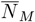 and distributions of mitochondria lengths, *L*_*M*_ obtained from the models M1 and M2 for the cells with experimental data (E) for MT_long_ cells. (A) Temporal evolution of 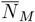. (B) Comparison of *L*_*M*_ . (C) Probability distribution, P(*L*_*M*_) of mitochondrial lengths in simulations with the Gaussian fits and comparison with experiments (D)

In experiments [1], this shift in the population toward longer mitochondria in MT_long_ cells is driven in part by the inability of the Dnm1 ring proteins to encircle mitochondria bound to the microtubules, thereby reducing the fission probabilities. We observe these deviations in the mitochondrial populations despite using the fission and fusion rates observed in the experiments. The situation is similar for both the M1 and M2 models. Since fission was assumed to be a first order process and fusion a second process, fission propensities were generally higher compared to the fusion propensities in these substantially low number density population systems. This could explain, in part, the increased tendency toward smaller mitochondria during the evolution of the dynamics in the KMC simulations. Furthermore, when microtubules are present, the modulation of the fission and fusion frequencies are only incorporated from a mean field perspective and the model is unable to predict the presence of the longer population of mitochondria observed in the experiments. In what follows we incorporate the presence of microtubules in a more realistic manner within the KMC framework by coupling the microtubule dynamics to the mitochondrial evolution within the cells.

### Mitochondrial population evolution in the presence of microtubules

The dynamics of microtubules were evolved using the Gillespie algorithm in the four sector model of the yeast cell (Fig 1C). Microtubule dynamics captured the changes in the microtubule lengths, controlled by polymerization, hydrolysis and depolymerization rates given by the rate constants *K*_*poly*_, *K*_*hy*_ and *K*_*depoly*_ respectively (Table 2). The optimized rate constants obtained using a genetic algorithm [57] reveals that the ratio of *K*_*hy*_ to *K*_*poly*_ is the highest for MT_short_ cells, implying higher rates of hydrolysis compared to rate of polymerization. However, in the case of MT_long_ the ratio of *K*_*hy*_ to *K*_*poly*_ was the lowest.

**Table 2.**
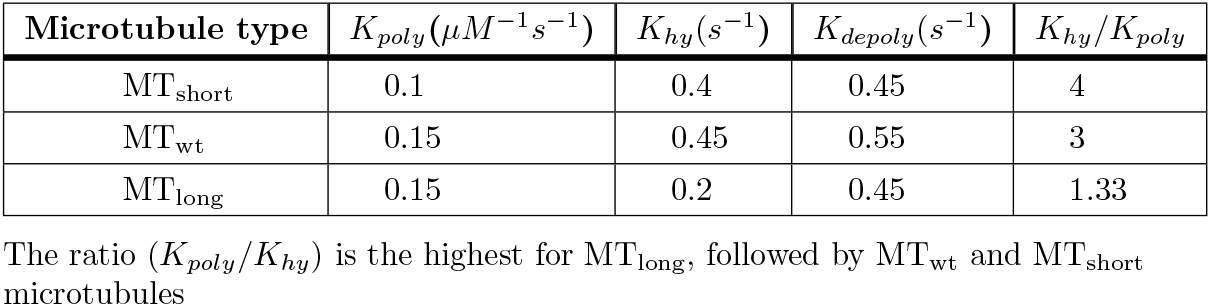
Rate constants for microtuble dynamics in cells with different microtubule environments.

| Microtubule type | $K_{poly}(\mu M^{-1} s^{-1})$ | $K_{hy}(s^{-1})$ | $K_{depoly}(s^{-1})$ | $K_{hy}/K_{poly}$ |
| --- | --- | --- | --- | --- |
| MT<sub>short</sub> | 0.1 | 0.4 | 0.45 | 4 |
| MT<sub>wt</sub> | 0.15 | 0.45 | 0.55 | 3 |
| MT<sub>long</sub> | 0.15 | 0.2 | 0.45 | 1.33 |

The ratio ( $K_{poly}/K_{hy}$ ) is the highest for MT<sub>long</sub>, followed by MT<sub>wt</sub> and MT<sub>short</sub> microtubules

For each of the different microtubule environments 1000 replicates were carried out and each simulation was evolved for at least 1000 s. The time evolution of the microtubule lengths, *L*_*MT*_ are illustrated in Fig 6 for a given replicate. The highest number of catastrophe events were observed for the MT_short_ cells (Fig 6A) followed by the MT_wt_ cells (Fig 6B), with the MT_long_ cells (Fig 6C) exhibiting the least number of catastrophe events. During a catastrophe event the microtubule length monotonically reduces to zero. As a result, the mean length for the MT_short_ cells was less than 40 % of the sector length (Fig 1A) whereas the mean length for MT_wt_ was ∼ 60 % and for the MT_long_ cells was ∼ 80 % of the sector length. For each of the different microtubule environments, parameters that characterize the microtubule dynamics such as elongation velocity 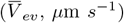, shrinkage velocity 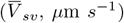 and catastrophe frequency 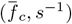 were computed by averaging across 1000 replicates for a given system and compared with the experiments carried out in yeast cells (Table 3). We observed that the elongation and shrinkage velocities for all the three different environments were accurately captured, however deviations were observed for the catastrophe frequencies for MT_short_ cells.

**Table 3.** Microtuble dynamics parameters.

| Microtubule type | $\bar{V}_{ev}(\mu m s^{-1})$ | | $\bar{V}_{sv}(\mu m s^{-1})$ | | $\bar{f}_c(s^{-1})$ | |
| --- | --- | --- | --- | --- | --- | --- |
| $MT_{short}^{**}$ | 0.0392 | 0.03** | 0.16 | 0.1** | 0.0015 | 0.02** |
| $MT_{wt}^*$ | 0.0543 | 0.0425* | 0.2 | 0.14* | 0.0015 | 0.0055* |
| $MT_{long}^*$ | 0.0542 | 0.05* | 0.075 | 0.1* | 0.0007 | 0.00067* |

\*Represent experimentally observed parameter values [58, 59]
\*\*Data procured from experiments carried out for current study

**Fig 6.**
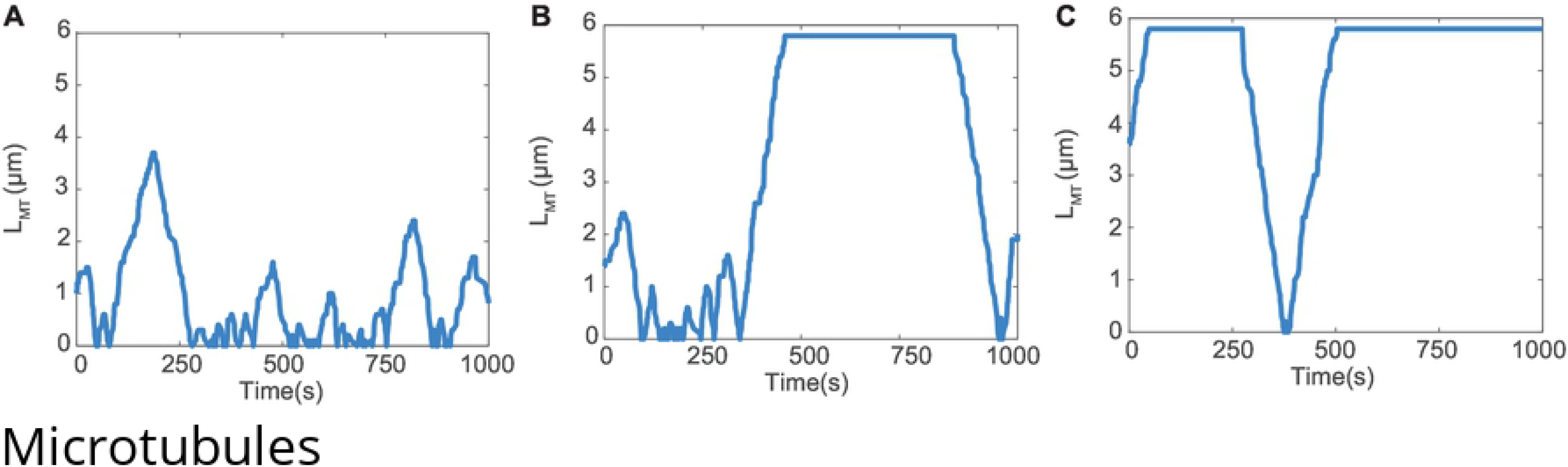
Microtubule temporal evolution dynamics (A) MT_short_ (B) MT_wt_ (C)MT_long_ environments. Here 5.8 *µ*m is the maximum length of a microtubule bundle which is also equal to the sector length.

In order to couple the microtubule dynamics to the mitochondrial evolution we carried out the following post processing steps on the microtubule lengths for each of the different cellular environments. Since mitochondria can grow to lengths greater than the horizontal sector or half the cell length (*L*_*s*_,Fig 1A) we superposed the microtubule lengths across two adjacent horizontal sectors at a given time to permit mitochondrial growth up to a maximum of the cell length (Fig 1D). We point out that 40 % of the mitochondrial lengths in the MT_wt_ and MT_long_ cells had lengths greater than *L*_*s*_ thereby justifying this superposition. Out of the 1000 replicates for a given environment we selected a single superposed length versus time evolution which was used as input into the KMC simulations for the mitochondrial evolution. The corresponding 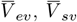 and 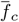 for this particular evolution was found to be close to the values reported in the Table 3 and are given in S6 Appendix. This selection was carried out by examining the length versus time data and selecting a replicate whose lengths were at least 80 % of the time within the maximum and minimum length bounds observed in the experiments. We also tested this criterion by varying the percentage of time that the microtubule lengths lie within the experimental bounds and we obtain similar number and lengths for the mitochondrial populations for microtubules lengths above 70%. However deviations were observed for the mitochondrial populations when microtubule lengths were below 70% (S6 Fig.).

The number and length distributions of mitochondria obtained after implementing the microtubule constraints on mitochondrial fission dynamics in the M2MT model are illustrated in Fig 7, Fig 8 and Fig 9. In this model, mitochondria undergo fusion reactions with their neighbors present in the same sector, similar to the previous M2 model as discussed in the Methods section. Fig 7A, 8A and 9A illustrates the mean number of mitochondria for the different microtubule cellular environments. For the MT_long_ cells (Fig 9A), the mean number decreased after the first time instant and remained saturated for about 400 s close to the experimentally observed value. This saturation in the mean number of mitochondria following an initial decrease is due to a higher frequency of fusion reactions when compared with fission reactions for the MT_long_ cells. On the contrary, in MT_short_ cells and MT_wt_ cells (Fig 7A and (Fig 8A, the mean number of mitochondria showed the greatest variation among the different cases. This was also reflected in the microtubule dynamics where a larger number of growth, shrinkage and catastrophe events occur (Fig 6). Such trends are caused due to a greater degree of temporal variation associated with the lengths in the MT_short_ and MT_wt_ cells compared to the growth dynamics of the MT_long_ cells (Fig 6). In experiments, the for MT_wt_ cells, the mean number of mitochondria was nearly invariant, around four, as predicted from model M2MT (Fig 8A). Thus, unlike previous models M1 and M2, incorporating microtubule dynamics improved the predictions of the mean mitochondria numbers resulting from approximately equal fission and fusion frequencies as observed in experiments for the MT_wt_ cells. Interestingly, in MT_short_ cells, however, the model M2MT showed an initial decreasing trend followed by a two-fold rise in the mean number of mitochondria as observed in experiments.

**Fig 7.**
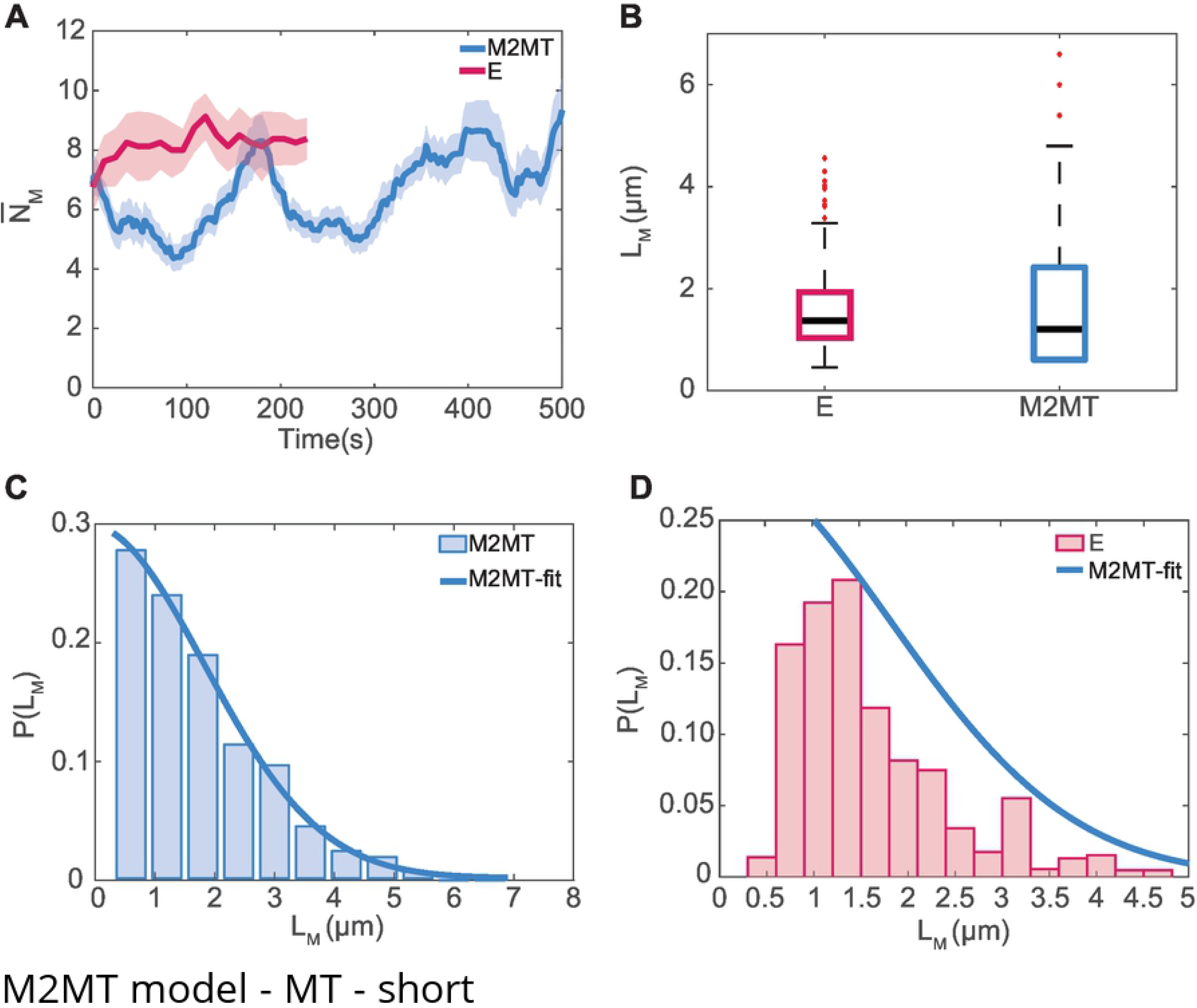
MT_short_ environment in M2MT model: (A)Temporal evolution trends of mean number of mitochondria (B) variation of mitochondrial lengths (C) Probability(P) distribution of mitochondrial lengths in experiments (D) Probability distribution and fit of mitochondrial lengths in M2MT model

**Fig 8.**
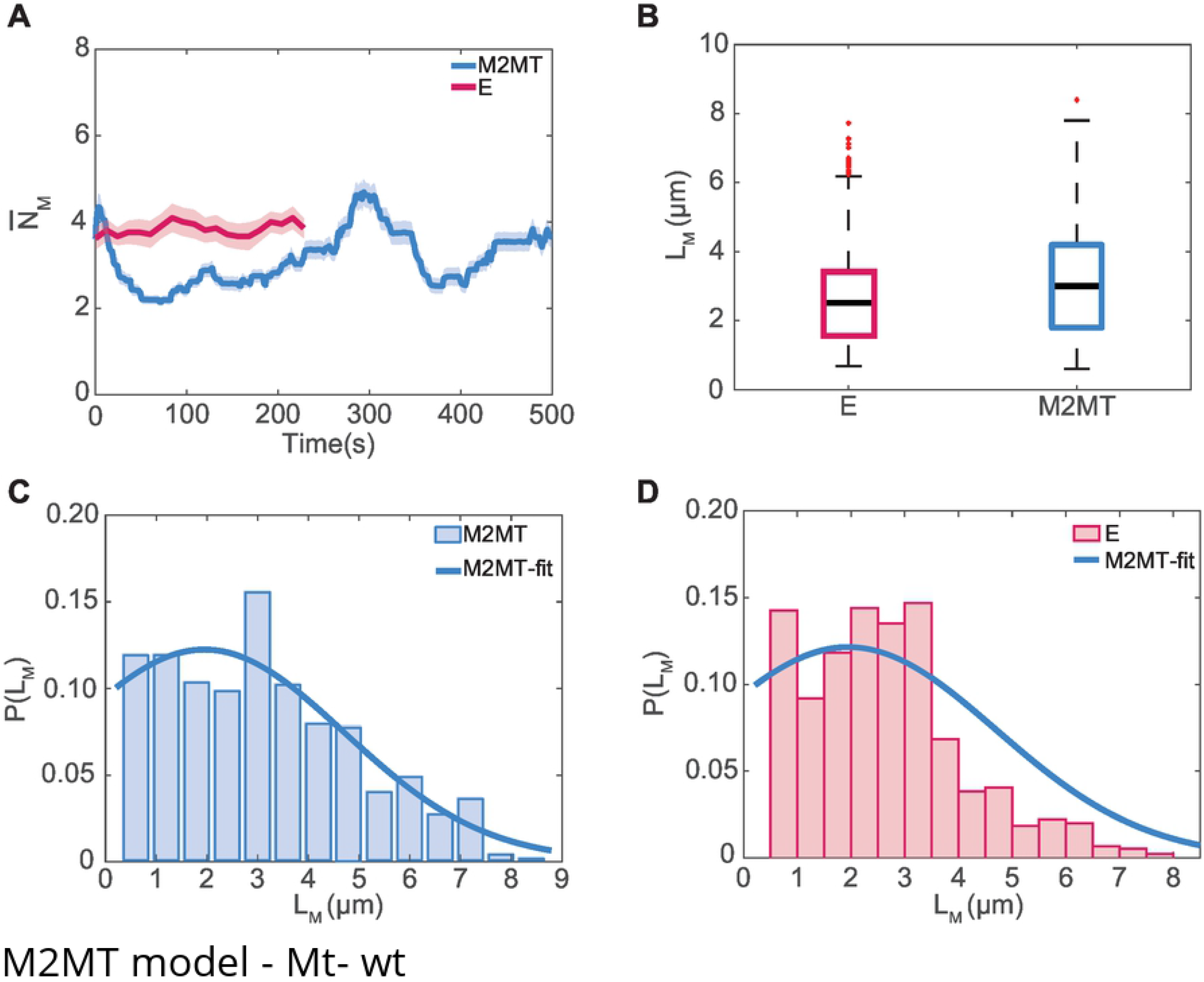
MT_wt_ environment in M2MT model: (A) Temporal evolution trends of mean number of mitochondria (B) variation of mitochondrial lengths (C) Probability (P) distribution of mitochondrial lengths in experiments (D) Probability distribution and fit of mitochondrial lengths in M2MT model

**Fig 9.**
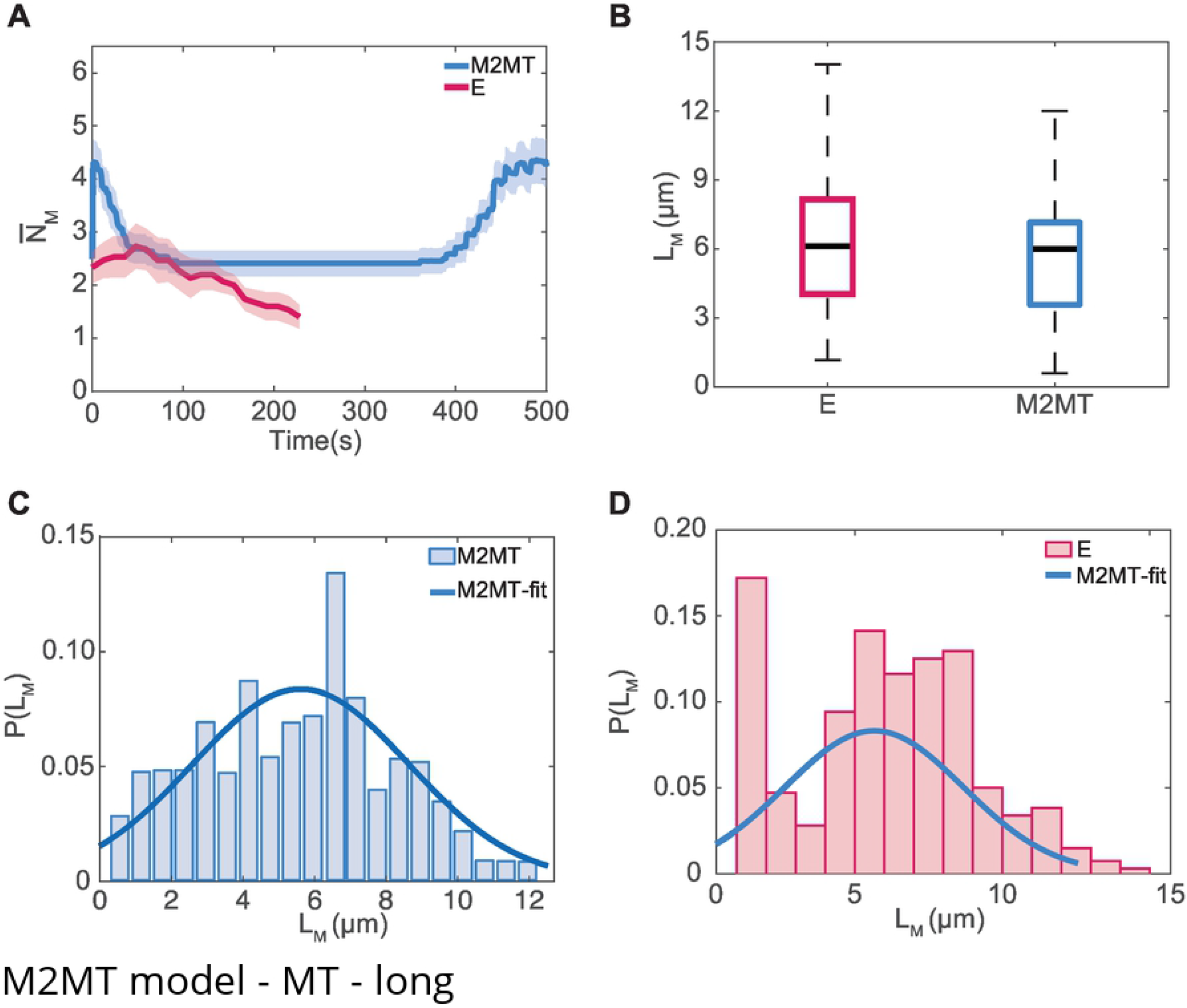
MT_long_ environment in M2MT model: (A) Temporal evolution trends of mean number of mitochondria (B) variation of mitochondrial lengths (C) Probability(P) distribution of mitochondrial lengths in experiments (D) Probability distribution and fit of mitochondrial lengths in M2MT model

However, the length distributions (Fig 7B, Fig 8B and Fig 9B) of mitochondria predicted by the model M2MT in all the three microtubule environments captured the experimental median values and the corresponding spread of the length distribution. Furthermore, the median length was the highest in MT_long_, followed by the MT_wild_ and MT_short_. The probability of lengths of mitochondria predicted from M2MT model in MT_short_ cells was peaked around the shortest mitochondria length of 0.6 *µ*m as illustrated in (Fig 7D). The Gaussian fit obtained from the M2MT model predicted the observed mitochondrial length probabilities in MT_short_ cells (Fig 7C) for lengths longer than 1.2 *µ*m, which was slightly higher margin of prediction of observed probabilities compared to that of 1.8 *µ*m in M1 and M2 model. Moreover, in MT_wt_ cells, the probabilities of mitochondrial lengths is peaked around 3.5 *µ*m (Fig 8D) similar to that observed in experiments (Fig 8C). Coupling of microtubule dynamics to that of mitochondria, shifted the peak from 0.6 *µ*m (Fig 4C) to 3.5 *µ*m. Similarly, for MT_long_ cells, the probability of mitochondrial lengths predicted from M2MT model is peaked at 7 *µ*m (Fig 9D), consistent with the experiments (Fig 9C). The Gaussian fit obtained from the M2MT model accurately predicted the probabilities of experimentally observed mitochondrial lengths for both MT_wt_ and MT_long_ cells(Fig 8C and Fig 9C). Moreover, in previous M1 and M2 models, irrespective of microtubule environments, since fission propensities were inherently high compared to that of fusion, a higher population of shorter mitochondria was observed (Fig 4B and 5B). Thus, by coupling microtubule dynamics to the mitochondrial dynamics, the shortcomings of models M1 and M2 were overcome in the current model M2MT.

## Conclusion

Modeling the evolution of the mitochondrial population in fission yeast cells is challenging due to the small number of mitochondria present in the cells and the high degree of stochasticity associated with their dynamics, as observed in experiments. Although the microtubule environment in known to play a critical role in mitochondria fission and fusion dynamics, [1] the coupling between the mitochondria and microtubule dynamics have not been reported in the literature. In this work, we use KMC simulations to predict the temporal evolution of mitochondria in both the mutated and wild-type states of microtubules in fission yeast cells. The mitochondrial population evolves through a reaction network involving fission and fusion reactions among the different mitochondrial species. Several models with varying complexity have been developed, and predictions of the mitochondrial populations have been compared with experimental data on fission yeast cells [1]. In the first class of models (M1 and M2), the fission yeast cell was treated as a well-mixed system, and mitochondrial fission and fusion rate constants were obtained from experiments. Interestingly when compared with the M1 model, the predictions from the M2 model where the cell had a single partition were in better agreement with experimental data. However, both models M1 and M2 predicted a higher mean number of mitochondria and shorter lengths of mitochondria compared to the experiments for MT_wt_ cells and MT_long_ cells. The deviation between the predicted values and the observed data from experiments is due to the higher frequency of first-order fission reactions when compared with the fusion reactions, which are second-order, in Mt_wt_cells and MT_long_ cells. Moreover, in experiments for Mt_wt_cells and MT_long_ cells, the shift in the population toward the lower mean number of mitochondria and longer mitochondria lengths was driven partly by the inability of the Dnm1 ring proteins to encircle mitochondria bound to the microtubules and the subsequent reduction in the fission probabilities.

The limitations of the M1 and M2 models in predicting the mitochondrial number and size distributions were overcome in the M2MT model where the microtubule and mitochondrial dynamics were coupled. Independent KMC simulations to predict the microtubule dynamics were carried out to obtain microtubule parameters such as elongation velocity, shrinkage velocity, and catastrophe frequency that were in good agreement with the experiments. Additionally, the median and spread of the mitochondrial length distributions were also captured quite accurately in the M2MT model. Our study reveals that the temporal evolution of mitochondrial populations is an intrinsic function of the presence of microtubules which modulates the fission and fusion frequencies to maintain mitochondrial homeostasis within cells. Furthermore, to realistically capture the mitochondria dynamics, it was essential to couple the microtubule polymerization and depolymerization dynamics with the kinetic Monte Carlo framework. An interesting application of the model lies in predicting the mitochondrial number distribution and morphology under different external stimuli such as cell division, temperature, and pH and capture changes in the mitochondrial distributions in response to these stimuli. The simulation framework developed in this work could potentially lead to a broader knowledge of the nature of mitochondrial dynamics in cells under different environments of stress and the underlying connection with disease states.

## Acknowledgments

We thank the Supercomputer Education and Research Center (SERC) computational facility at the Indian Institute of Science, Bengaluru, for access to supercomputing resources. K.G.A. acknowledges funding by a grant from the Department of Science and Technology, Government of India. We also thank Sanjeev K Gupta, Danny Raj M and Ashwin Nair for several useful discussions on the simulations reported in this manuscript.

---

**Supplementary figure.**
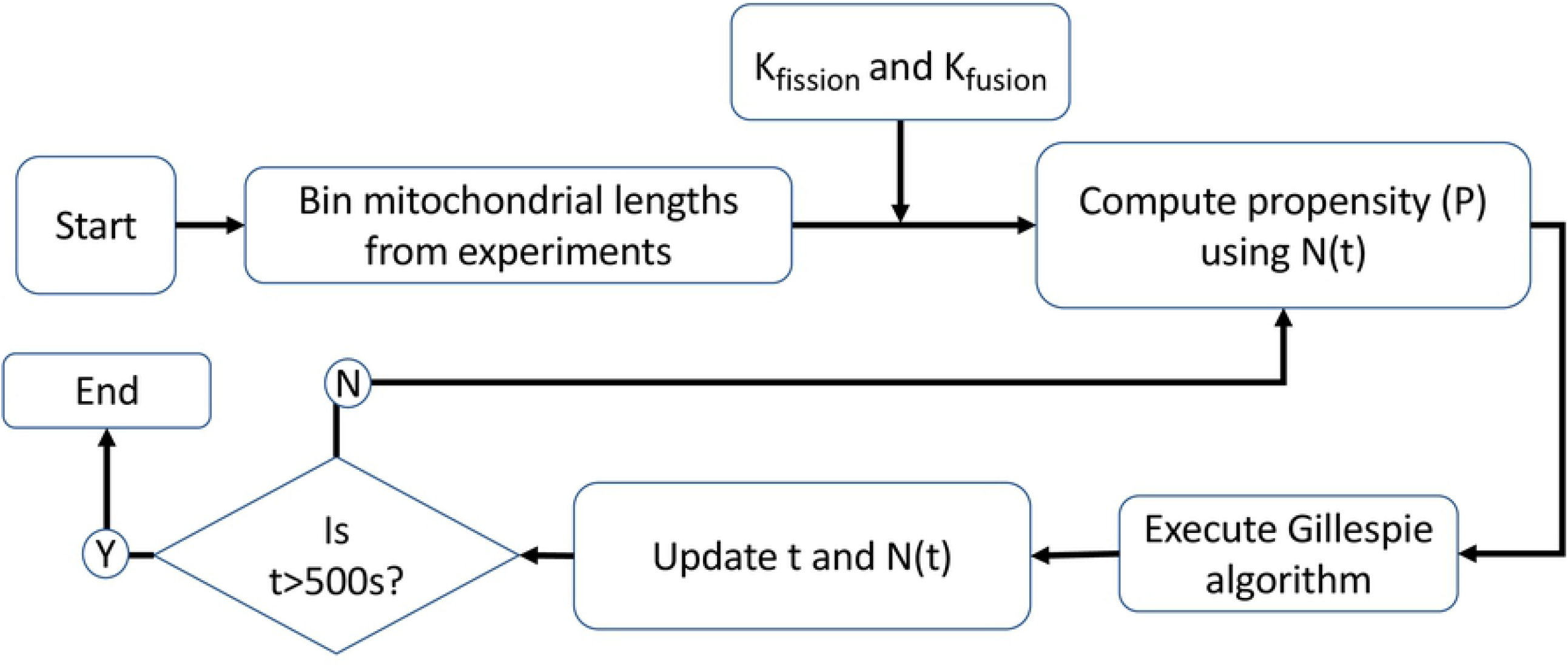
M1-model algorithm

**Supplementary figure.**
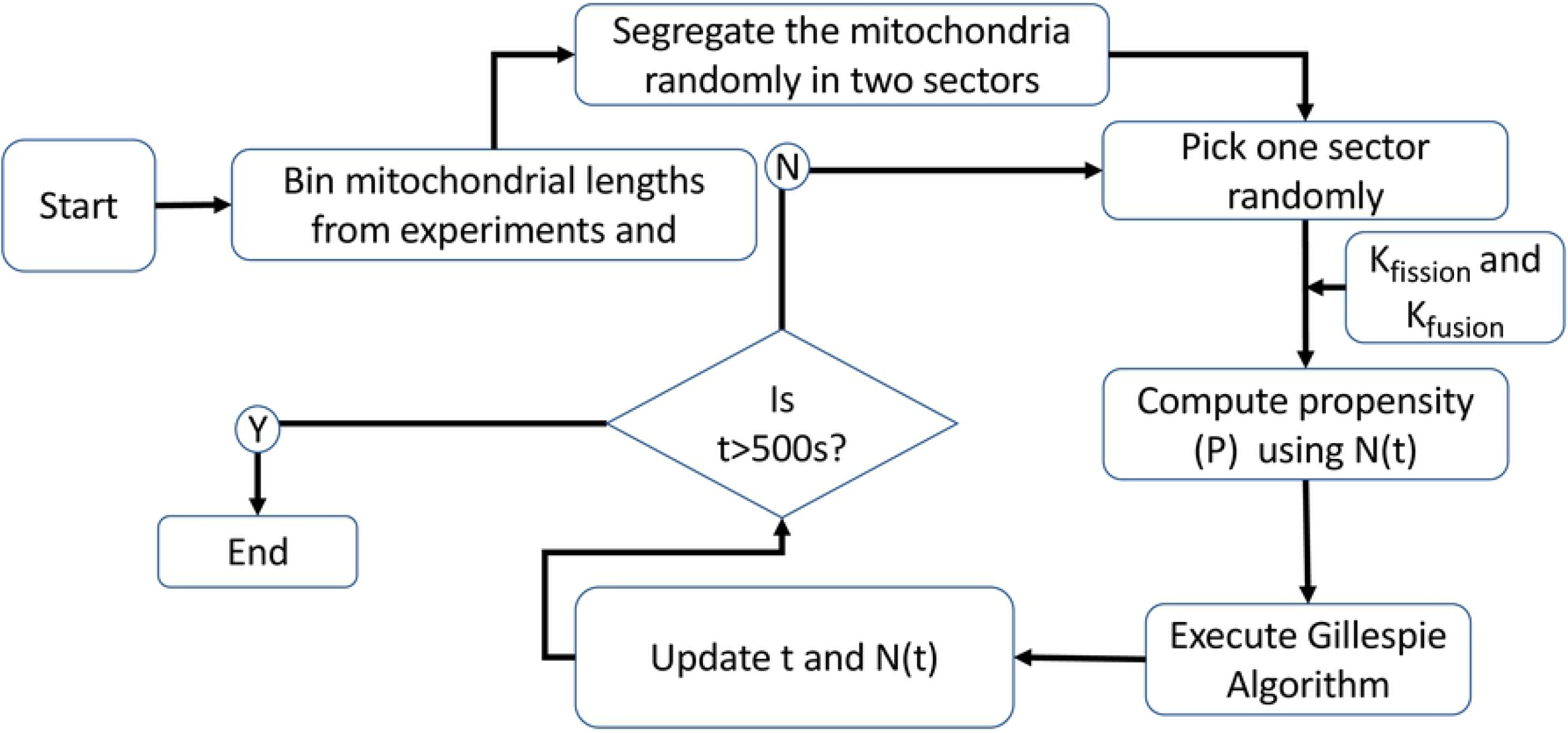
M2-model algorithm

**Supplementary figure.**
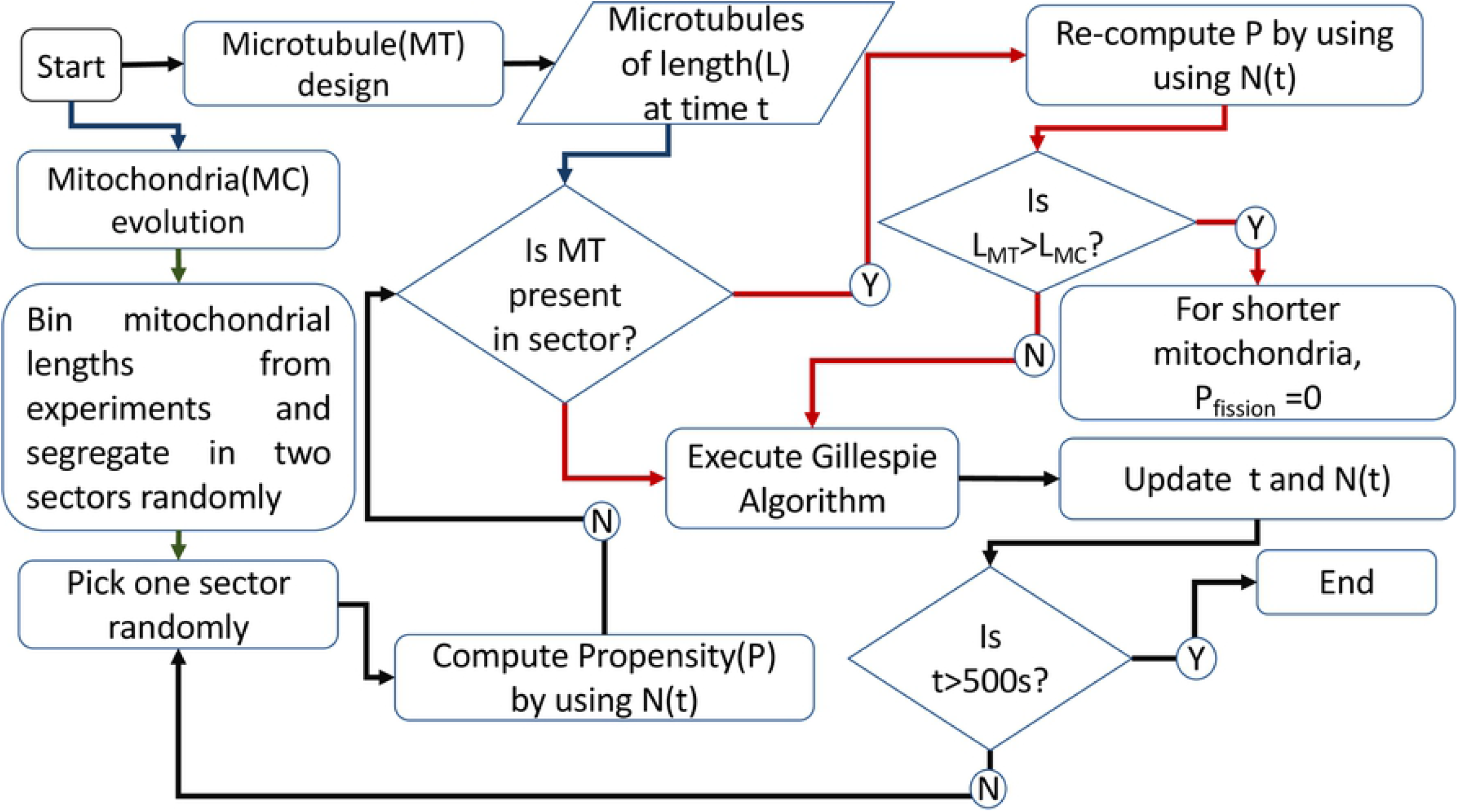
M2MT-model algorithm

**Supplementary figure.**
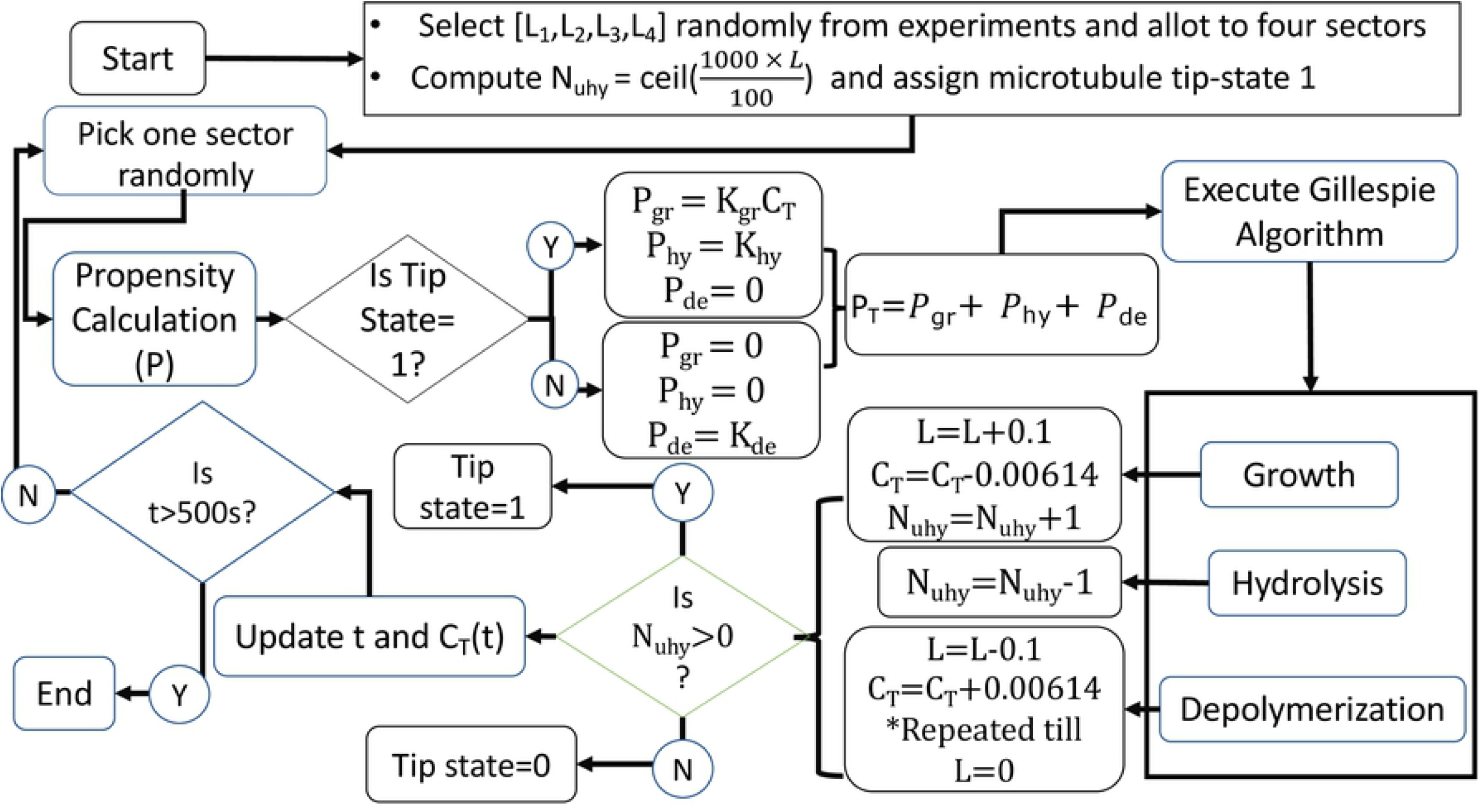
Microtubule design algorithm

**Supplementary figure.**
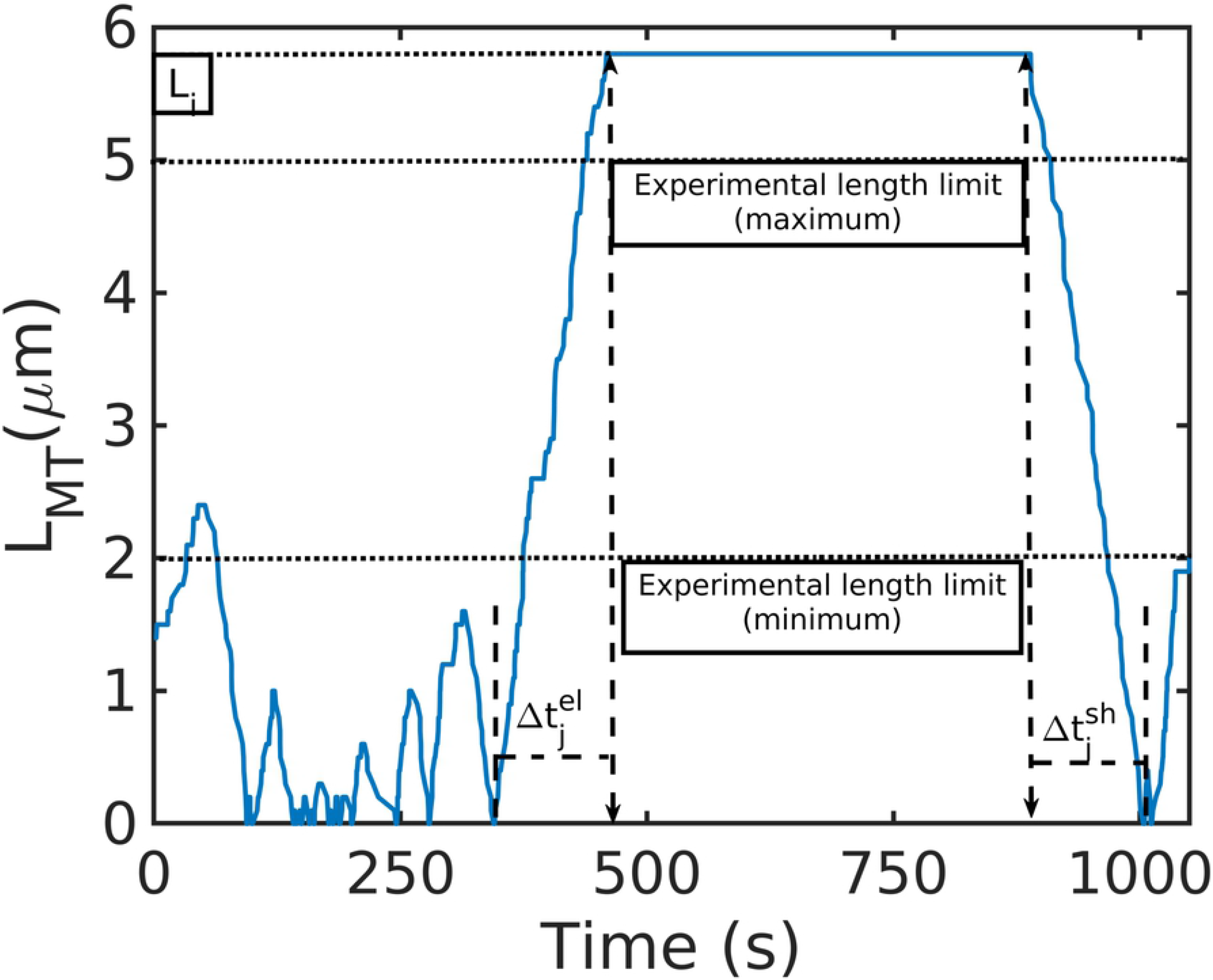
Microtubule parameters

**Supplementary figure.**
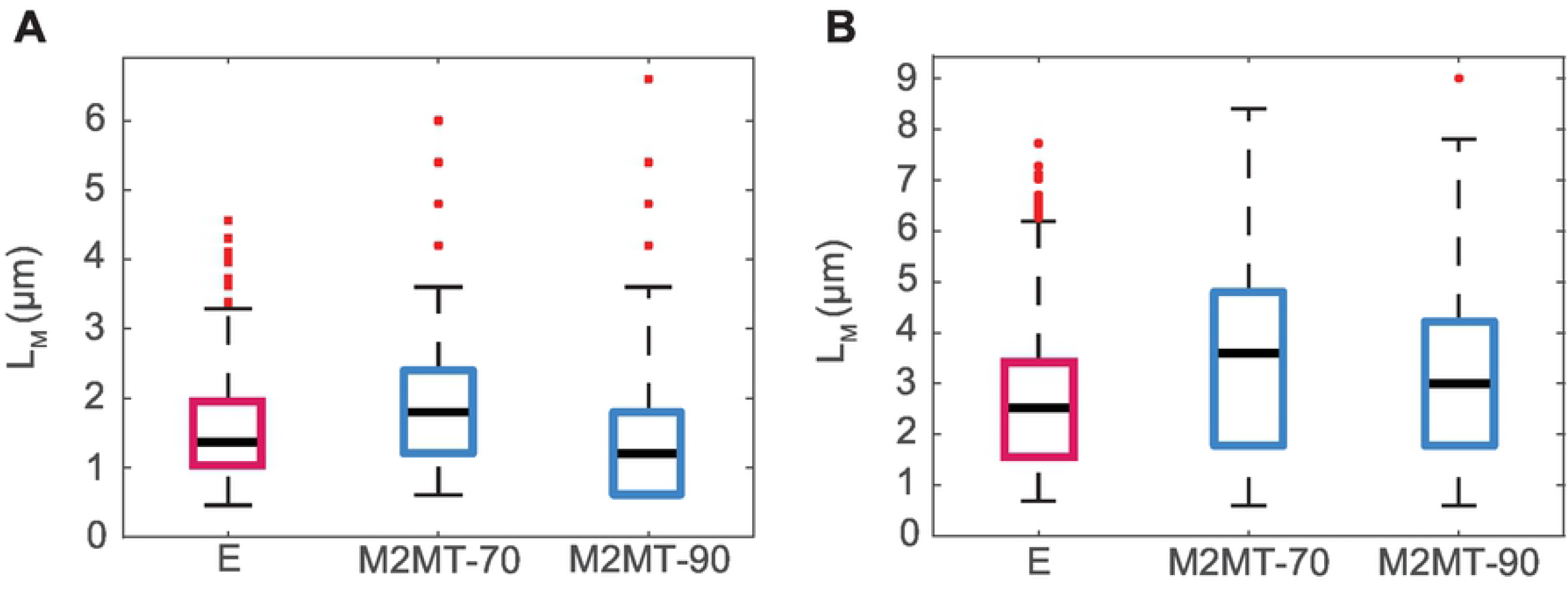
Microtubule occupancy

